# Efficient generation of endogenous protein reporters for mouse preimplantation embryos

**DOI:** 10.1101/2020.08.27.266627

**Authors:** Dan O’Hagan, Amy Ralston

## Abstract

Fluorescent proteins and epitope tags can reveal protein localization in cells and animals. However, the large size of many tags hinders efficient genome targeting. Accordingly, many studies have relied on characterizing overexpressed proteins, which might not recapitulate endogenous protein activities. We present two approaches for higher throughput production of endogenous protein reporters. Our first approach makes use of a split fluorescent protein mNeonGreen2 (mNG2). Knock-in of a small portion of the *mNG2* gene, in frame with gene coding regions of interest was highly efficient in embryos, eliminating the need to establish mouse lines. When complemented by the larger portion of the *mNG2* gene, fluorescence was reconstituted and endogenous protein localization faithfully reported in living embryos. However, we report a threshold of detection using this approach. By contrast, the V5 epitope enabled high efficiency and higher sensitivity protein reporting. We describe complementary advantages and prospective applications of these two approaches.

**Highlights:**

- Split fluorescent protein for in vivo protein localization in living embryos
- V5 tagging for in vivo localization of low abundance proteins
- Bypassing the need for founder mouse lines for preimplantation studies
- Guidelines and strategies for implementation and prospective applications

## Introduction

Mouse preimplantation embryos have long served as a paradigm for discovering the molecular regulation of cell fate specification. This is in part because preimplantation embryos provide certain experimental advantages, including optical transparency, the capacity to develop ex vivo in a cell culture incubator, robust tools for genome editing, and facility of collecting dozens of embryos at desired developmental time points. Additionally, preimplantation development is an alluring model for scholarly reasons. During these developmental stages, the first two lineage decisions establish three distinct cell types that are all, in fact, stem cell progenitors (Frum and Ralston, 2015; Posfai et al., 2014; White and Plachta, 2020). Therefore, mouse early development enables dissection of molecular regulation of stem cell origins in a biologically meaningful, yet relatively simple, setting.

One powerful approach to elucidating the molecular mechanisms of development has been live imaging of fluorescent reporters, which enables analysis of fate decisions at the single cell level (Nowotschin and Hadjantonakis, 2014). Live imaging of protein localization in the embryo is often achieved by injection of exogenous mRNAs encoding fluorescently tagged proteins. While efforts are made to ensure that injection does not cause overt overexpression phenotypes, imaging endogenous proteins could provide certain advantages. Genes encoding Green Fluorescent Protein (GFP) and other fluorescent proteins have been knocked into loci to enable visualization of endogenous protein and gene expression patterns. However, this approach is not as common as it could be, likely due to lengthy breeding required to establish new mouse lines.

We have explored the use of two alternative approaches for tagging endogenous proteins in vivo. The first system takes advantage of the observation that fluorescent proteins such as GFP can be split into large and small coding units that, when coexpressed, will produce two proteins capable of self-assembly through non-covalent intermolecular interactions and reconstitution of fluorescence (Cabantous et al., 2005). This approach has been used in cell lines (Feng et al., 2017; Kamiyama et al., 2016; Leonetti et al., 2016), where a larger portion of the fluorescent protein-encoding gene is introduced virally, and is then complemented by knocking in the smaller portion of the fluorescent protein into loci of interest, in frame with the coding region. While effective in cultured cells, this approach has not been tested in an animal model, where it could provide the most insight into biologically relevant processes. Here, we report that split mNeonGreen2 (mNG2) can be used in mice to visualize the localization patterns of essential proteins in living embryos.

Our second approach makes use of the V5 epitope, commonly used for antibody-based purification of overexpressed proteins in cell lines for applications such as chromatin immunoprecipitation and sequencing (ChIP-seq). V5 has been knocked into endogenous mouse loci (Dewari et al., 2018; Yang et al., 2013), but its sensitivity in terms of protein detection, compared to fluorescent fusion proteins, is unreported. We show that targeting efficiencies for both the split fluorescent protein the V5 systems are sufficiently high so as to eliminate the need to establish new mouse lines prior to evaluating protein expression patterns during early embryogenesis. We compare the sensitivities of these two systems to provide guidelines for their optimal implementation. Finally, we discuss applications of these two systems for discovery of novel regulatory mechanisms of development.

## Design

Our goal was to develop a pipeline for detection of endogenous proteins in mice. We sought to overcome challenges associated with knocking in large fluorescent protein tags and with screening commercially available antibodies, including cost, time, and effort. We therefore selected two relatively small (14-16 amino acid) tags to enable more efficient knock-in in mouse embryos. Knock-in targeting constructs were synthesized commercially as a single stranded oligonucleotide nucleotide (ssODN) donors by commercial vendors, further streamlining the pipeline. Each of these tags provides complementary advantages for in vivo imaging of endogenous proteins. The first approach makes live imaging of endogenous proteins more efficient by splitting the fluorescent protein in two. The second makes detection of diverse, low abundance proteins more straightforward by making use of high signal to noise, commercially available reagents.

## Results

### A mouse line to enable in vivo implementation of a split fluorescent protein

The yellow-green, monomeric fluorescent protein mNeonGreen, derived originally from the marine invertebrate *Branchiostoma lanceolatum* is up to three times brighter than GFP (Shaner et al., 2013). Its derivative, mNG2 is can be split into two separate coding units, mNG2(Δ11), which lacks the eleventh beta barre, and mNG2(11) (Feng et al., 2017). Individually, the two resulting proteins lack appreciable fluorescence. However, when the larger protein mNG2(Δ11) is complemented by the 16-amino acid mNG2(11), fluorescence is reconstituted (Fig. 1A).

**Figure 1.**
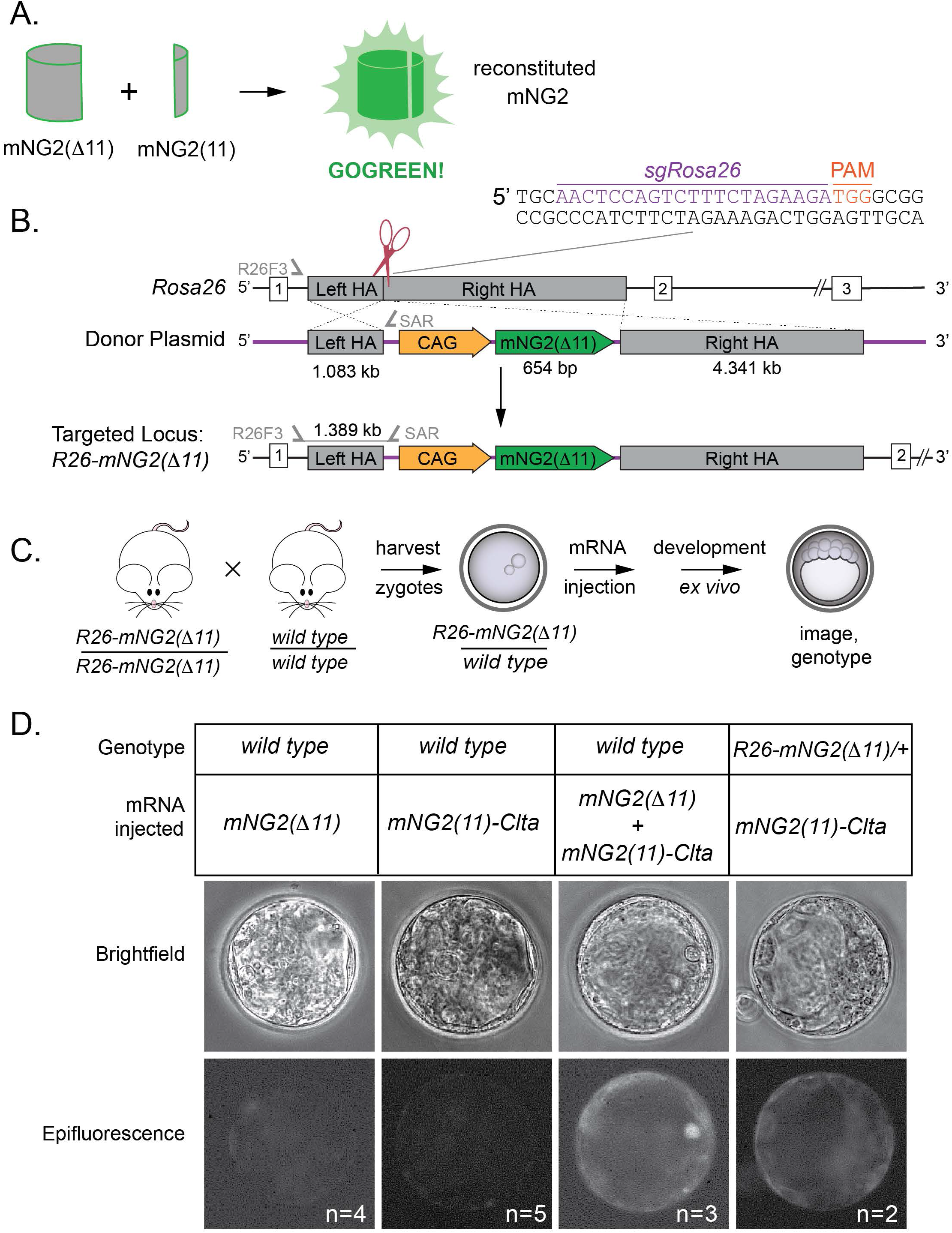
Development of the GOGREEN mouse line. A) Deletion of the eleventh beta-barrel (Δ11) of the fluorescent protein mNeonGreen2 (mNG2) eliminates its fluorescent properties. However, complementation by coexpression of mNG2(Δ11) and the eleventh beta barrel mNG2(11) enables non-covalent association of the two proteins and reconstitution of the mNG2 fluorescent properties. B) Strategy for CRISPR/Cas9 knock in of the *mNG2(Δ11)* expression construct into the mouse *Rosa26* locus. Sequence of single guide RNA (sgRNA), location of genotyping primers (R26F3 and SAR) and predicted Cas9 cut site are shown. HA = homology arm. C) Genetic strategy for testing fluorescent complementation in early embryos. Zygotes carrying *mNG2(Δ11)* were harvested, and then injected with mRNA encoding mNG2(11)-tagged Clathrin (CLTA). Zygotes were subsequently cultured ex vivo to later stages and fluorescence examined in individual embryos. Individual embryo genotypes were determined by PCR. In control experiments, zygotes were produced from wild type parents. D) Evidence of fluorescent reconstitution in vivo using the GOGREEN system. Negative controls: wild type embryos injected with mRNA encoding only *mNG2(Δ11)* or *mNG2(11)*-tagged *Clta* exhibit background levels of fluorescence (columns 1-2). Positive control: wild type embryos coinjected with *mNG2(Δ11)* and *mNG2(11)*-*Clta* exhibit reconstituted fluorescence (column 3). Test of *R26-mNG2(Δ11)* mice: knock-in embryos injected with mRNA encoding *R26-mNG2(11)-Clta* also exhibit reconstituted fluorescence above background, although level is apparently dimmer, likely owing to expression from the endogenous locus (column 4). n = number of embryos evaluated in this experiment.

We sought to make use of this fluorescence complementation strategy to evaluate localization of endogenous proteins in mouse embryos because we reasoned that tagging endogenous proteins with the relatively small *mNG2(11)* coding region would be more efficient than knocking in the full-length gene encoding the full-length fluorescent protein. Then, to provide the complementary protein, we aimed to establish a mouse line capable of constitutive expression of *mNG2(Δ11)*. Our goal was to introduce an expression construct including cytomegalovirus enhancer, chicken beta-actin promoter (CAG) and *mNG2(Δ11)* into the *Rosa26* (*R26*) locus by homologous recombination (Fig. 1B), which would enable constitutive, ubiquitous expression of *mNG2(Δ11)* throughout mouse tissues and development (Friedrich and Soriano, 1991). However, prior to attempting knock-in in mouse zygotes, we first established *R26-mNG2(Δ11)* embryonic stem (ES) cell lines using a Cas9/CRISPR-mediated knock-in strategy based on a previously published study (Chu et al., 2016) (see Methods). The *R26-mNG2(Δ11)* ES cells provided a renewable source of positive control genomic DNA for subsequent experiments.

To produce a mouse line capable of expressing *mNG2(Δ11)*, we introduced *CAG>mNG2(Δ11)* into the *Rosa26* (*R26)* locus in zygotes, following the strategy we had used in ES cells. Injected zygotes were transferred to recipient females, allowed to gestate, and then founder mice carrying *mNG2(Δ11)* were identified by PCR genotyping (Fig. S1A, B) and genomic sequencing (not shown). A single founder mouse was then expanded (Fig. S1C), and bred to homozygosity to establish the *R26-mNG2(Δ11)*/*R26-mNG2(Δ11)* mice. In principle, providing *mNG2(11)* in trans to *R26-mNG2(Δ11)* would lead to reconstitution of the green fluorescent protein. For simplicity, we called this the GOGREEN system.

To test the performance of the GOGREEN system in embryos, we first provided exogenous *mNG2(11)* by mRNA injection into *R26-mNG2(Δ11)*/+ zygotes (Fig. 1C). For negative controls, wild-type zygotes were injected with mRNAs encoding either *Clta-mNG2(11)* or *mNG2(Δ11)*. These embryos exhibited minimal fluorescence at blastocyst stage (Fig. 1D). For a positive control, wild type zygotes were co-injected with mRNAs encoding both *R26-mNG2(Δ11)* and *mNG2(Δ11)*, and these exhibited greatly elevated fluorescence at the blastocyst stage. Finally, *R26-mNG2(Δ11)*/+ zygotes were injected with mRNA encoding *Clta-mNG2(11)*, which led to elevated fluorescence at the blastocyst stage. These observations demonstrated functionality of the GOGREEN system using exogenous *mNG2(11)*.

### Fluorescence reconstitution by split fluorescent knock-in

We next aimed to evaluate the performance of the GOGREEN system when *mNG2(11)* was endogenously expressed. Our goal was to derive *R26-mNG2(Δ11)*/+ zygotes and in these, perform CRISPR/Cas9 knock-in of *mNG2(11)* (Fig. 2A) to produce mNG2(11) fusion proteins capable of complementing mNG2(Δ11) and reporting endogenous protein patterns.

**Figure 2.**
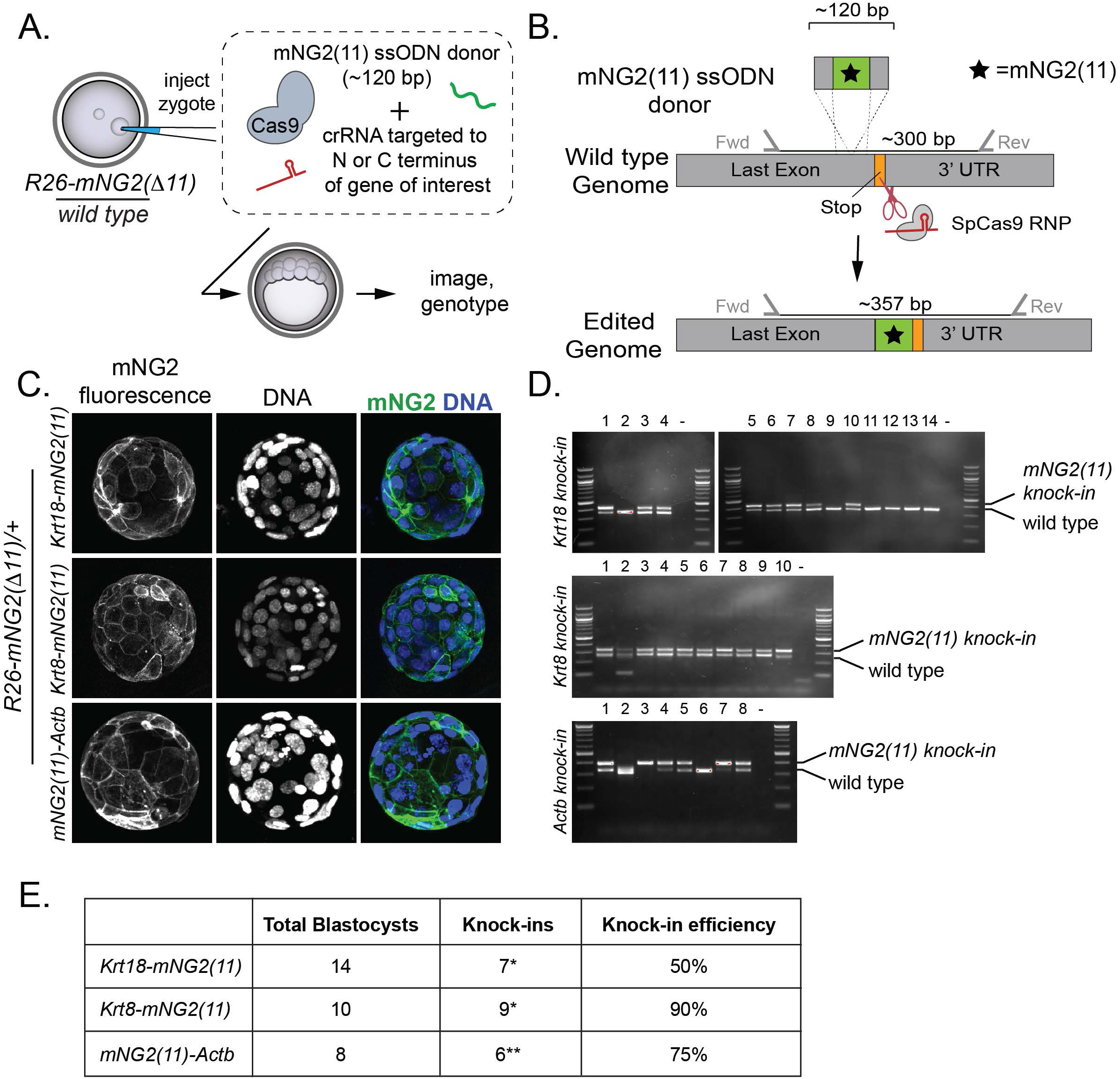
The GOGREEN system enables efficient production of reporters of several endogenous proteins in vivo. A) Experimental design: *R26-mGN2(Δ11)/+* knock-in zygotes (generated *per* cross shown in Fig. 1C) are injected with CRISPR/Cas9 targeting reagents to knock *mNG2(11)* into loci of interest, in-frame with encoded proteins. Embryos are then cultured ex vivo, imaged, and genotyped to evaluate the efficiency of mNG2(11) knock in. ssODN = single-stranded oligodeoxyribonucleotide, gRNA = guide RNA. B) Overview of strategy for targeting *mNG2(11)* to genomic loci to produce fusion proteins. C) The GOGREEN system enables detection of endogenous cytoskeletal proteins including intermediate filaments KERATIN18, KERATIN8, and ACTIN (ACTB). Note that 100% of embryos inherited *R26-mNG2(Δ11)*. D) PCR genotyping of embryos, including those shown in panel C, to identify which embryos were successful *mNG2(11)* knock-ins. *one additional embryo presented the knock-in genotype, but not the expected fluorescent phenotype. **one additional embryo presented the fluorescent phenotype but not the expected genotype. E) Summary of mNG2(11) knock-in results observed for the three loci shown.

To achieve in-frame *mNG2(11)* knock in, we designed targeting constructs encoding the 16 amino acid mNG2(11), plus a three-amino acid linker, flanked by genomic locus-specific homology arms of 30 nucleotides each (Fig. 2B). Resulting targeting constructs ranged from 117-120 nucleotides in length and could therefore be synthesized as a single stranded oligodeoxyribonucleotides (ssODN) by a commercial vendor. For our first knock-in attempts, we targeted cytoskeletal proteins, including intermediate filaments and beta-actin, because their subcellular localizations in mouse preimplantation have long been known (Chisholm and Houliston, 1987; Coonen et al., 1993; Reima and Lehtonen, 1985).

We designed CRISPR reagents to knock *mNG2(11)* in-frame with *Keratins* (*Krt*) *Krt8* and *Krt18*, as well as *Actin, beta* (*Actb)*. Following injection of the knock-in mixture into *R26-mNG2(Δ11)*/+ zygotes, embryos were cultured to the blastocyst stage, and then imaged by confocal microscopy. For all three genes, we observed the anticipated expression patterns of fluorescent fusion protein (Fig. 2C). Individual embryos were then harvested, and targeting evaluated by PCR (Fig. 2D) and sequencing (not shown). In all cases, targeting was highly efficient, as expected (Fig. 2E). These observations demonstrate the utility of the GOGREEN system for efficiently reporting localization of endogenous proteins in vivo.

### Fluorescence complementation in the nuclear compartment

Thus far, we had evaluated the ability of the GOGREEN system to report endogenous cytoplasmic proteins in vivo. Given that the dynamics of cytoskeletal protein localization and turnover during preimplantation development are actively studied (Anani et al., 2014; Schwarz et al., 2015; Zenker et al., 2018), the GOGREEN system could benefit the field. There has also been great interest in the live imaging of transcription factor dynamics in living preimplantation embryos (Gu et al., 2018; McDole et al., 2011; Posfai et al., 2017; Saiz et al., 2016). However, we were uncertain whether the GOGREEN system could effectively report nuclear proteins, owing to the possibility that the two components of the GOGREEN system might end up separated by the nuclear membrane.

To investigate the performance of the GOGREEN system for visualizing nuclear proteins in vivo, we surveyed fluorescence in embryos, after targeting diverse loci. As for previous experiments, we targeted *mNG2(11)* in frame with genes of interest in the *R26-mNG2(Δ11)*/+ genetic background. We were able to detect fluorescence within the nuclear compartment for some of the nuclear proteins surveyed (Fig. 3A), with confirmed knock-ins of *mNG2(11)* (Fig. 3B, C). However, for the majority of nuclear proteins surveyed, we were unable to detect fluorescence, in spite of apparently successful *mNG2(11)* knock-in (Fig. S2A), even though these proteins, including *Cdx2, Yap1, Gata6*, and *Nanog*, possess biologically essential activities during preimplantation (Frum et al., 2018; Mitsui et al., 2003; Schrode et al., 2014; Strumpf et al., 2005).

**Figure 3.**
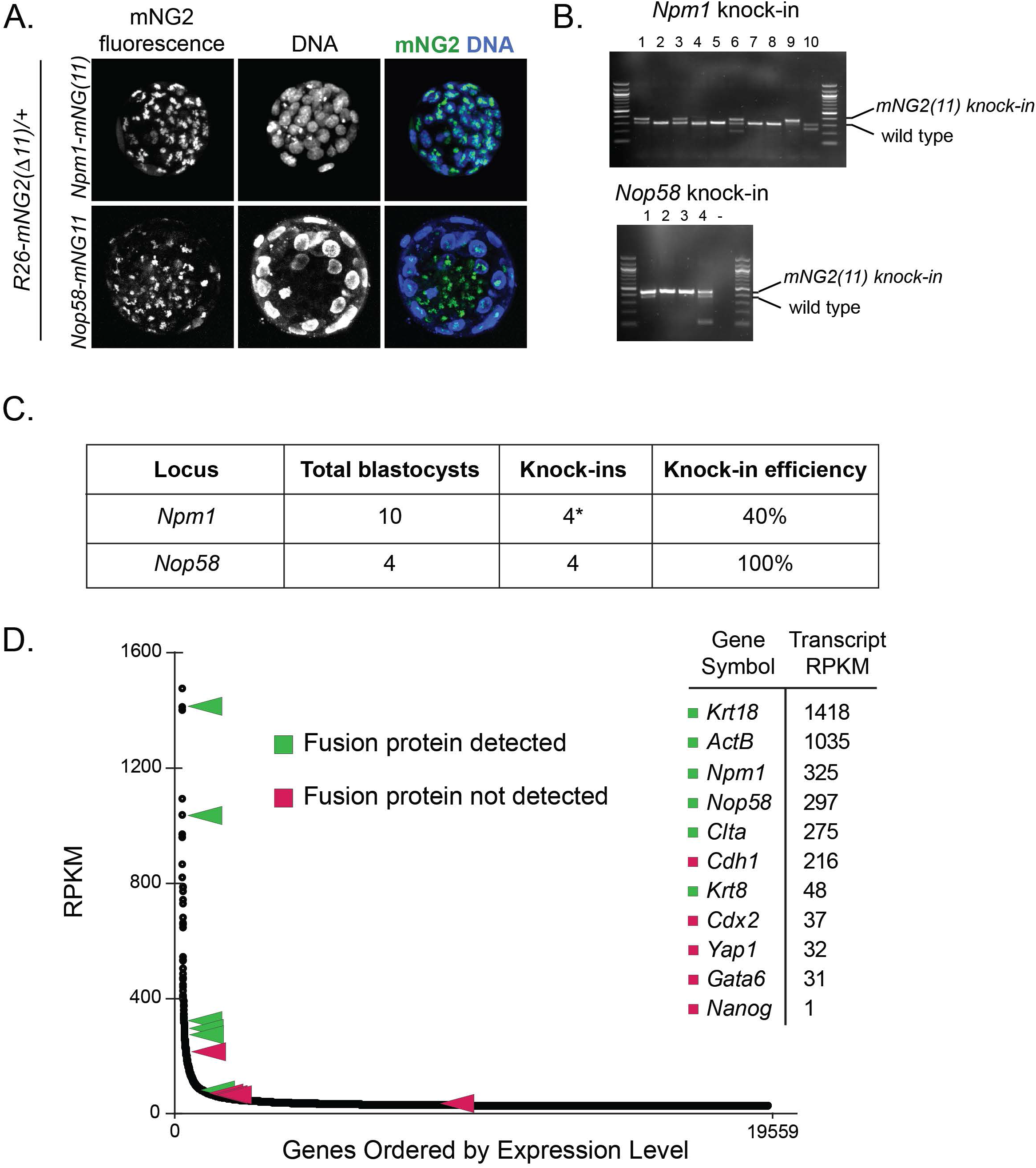
The GOGREEN system enables detection of abundant endogenous proteins in the nuclear compartment, but lacks sensitivity to detect transcription factors. A) In the *R26-mNG2(Δ11)* background, knock-in of *mNG2(11)* in-frame with the coding regions of two different nuclear proteins demonstrates fluorescent reconstitution of mNG2 and localization within nuclei. B) PCR genotyping to confirm knock-in of *mNG2(11)* into indicated loci for individual embryos, including those shown in panel A. *one additional embryo presented the fluorescent phenotype but not the expected genotype. C) Efficient knock-in of nuclear proteins shown in panels A-B. D) Relative abundance of endogenous mRNAs encoding tagged proteins evaluated using the GOGREEN system, as reported by RNA-seq analysis (Aksoy et al., 2013). Each black dot indicates a unique gene transcript, and RPKM = reads per kilobase per million reads. The GOGREEN system was able to detect relatively abundant nuclear proteins (green), but not less abundant nuclear proteins, including many transcription factors. For all genes shown, *mNG2(11)* knock-in was confirmed by PCR (Fig. S2).

Notably, our ability to detect specific proteins using the GOGREEN system correlated with the abundance of their transcripts in embryos (Fig. 3D). These observations suggest that, rather than subcellular localization, target protein abundance is the limiting factor for detection using the GOGREEN system. Consistent with this suggestion, we were unable to detect the relatively low abundance, non-nuclear protein E-cadherin (CDH1) in these assays. Moreover, we were able to detect the low abundance nuclear protein CDX2 when *Cdx2-mNG2(11)* was overexpressed (Fig. S2B,C), arguing that protein level, rather than localization, is crucial for efficacy of the GOGREEN system.

### Robust detection of low abundance endogenous proteins

Because were unable to detect relatively low abundance, yet biologically important, factors in preimplantation embryos using the GOGREEN system, we sought to establish an alternative knock-in tagging strategy for detecting this class of proteins. In contrast to the GOGREEN system, epitope tagging does not enable live imaging, owing to tissue fixation during immunolocalization protocols. However, we reasoned that being able to image diverse proteins with a single antibody, of renewable source, would provide considerable advantages over identifying distinct antibodies recognizing individual proteins.

We selected the V5 epitope, a 14 amino acid region of proteins encoded in genome of paramyxovirus, a member of the simian virus 5 (SV5) family, because its small size promised efficient knock-in (Fig. 4A), and because of the existence of low background, commercially available, monoclonal anti-V5 antibody, which could be used for immunofluorescent detection of V5-tagged proteins in embryos. We then designed V5-encoding targeting constructs for generating in-frame V5 fusion proteins (Fig. 4B).

**Figure 4.**
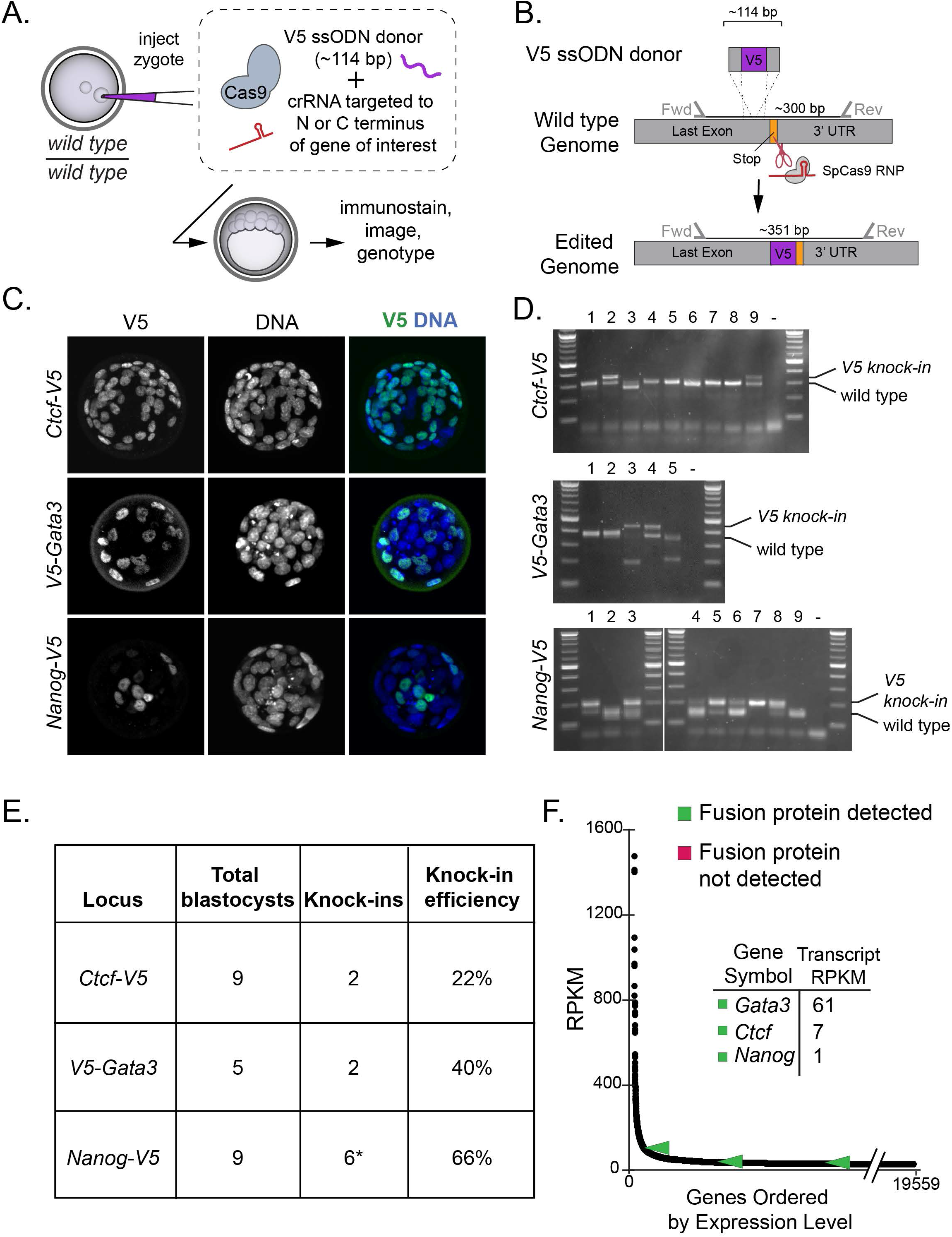
An alternative, V5-based system for detection of low abundance proteins *in vivo*. A) Strategy for knocking the V5-encoding gene into loci of interest in wild type zygotes, to enable detection of diverse endogenous proteins with an anti-V5 antibody. B) Overview of V5 targeting strategy. C) Examples of low abundance proteins detected as V5 fusion proteins, following knock-in as illustrated in panels A-B. D) PCR genotyping of embryos to confirm V5 knock-in at indicated genomic loci, including those shown in panel C. *one additional embryo presented the knock-in genotype, but not the expected fluorescent phenotype. E) Summary of V5 knock-in efficiency at indicated loci. F) Relative abundance of transcripts of interest in wild type embryos (Aksoy et al., 2013).

We targeted the V5 tag in frame with several transcription factors in zygotes, and then observed immunofluorescence patterns in blastocysts by confocal microscopy. We were able to detect V5 signals that were clear and specific after targeting the nuclear factors such as GATA3, CTCF, and NANOG (Fig. 4C-E). Importantly, the patterns of V5-tagged GATA3 and NANOG recapitulated their reported expression patterns in blastocysts (Home et al., 2009; Ralston et al., 2010; Strumpf et al., 2005). The expression pattern of CTCF has not been previously reported in blastocysts, although its DNA binding sites have been characterized in this context (Hainer and Fazzio, 2019). We detected tagged CTCF all nuclei of all cells of the blastocyst, consistent with RNA-seq data (Aksoy et al., 2013). These observations highlight the utility of the V5-tagging system to discover novel protein expression patterns.

Remarkably, transcript levels of *Gata3, Ctcf*, and *Nanog* are all extremely low in whole embryos, compared with other transcripts (Fig. 4F, compare with Fig. 3D). This observation indicates that the V5 system is highly sensitive. Thus, the V5 knock-in system, together with the GOGREEN system provide a pipeline for the characterization of proteins with diverse functions, over a range of expression levels, all within a biologically relevant developmental setting.

## Discussion

The conventional approach to live imaging endogenous proteins in embryos relies on the genetic transmission of engineered loci. Per Mendelian rules of inheritance, 25 to 50% of embryos typically inherit the transgene or knock-in allele encoding the fluorescent protein. As an alternative, we have exploited the intrinsically high efficiency of CRISPR/Cas9-mediated homologous recombination to knock in very small protein tags at efficiencies ranging from 22-100%, with an average knock-in efficiency of 59% for over more than a dozen knock-in experiments undertaken here. This degree of knock-in efficiency, observed within the F0 generation, obviates the need to establish and breed knock-in reporter mouse lines for preimplantation studies of protein localization.

### Limitations

For highly abundant proteins, we were able to observe fluorescence using the GOGREEN system. However, we were unable to detect less abundant proteins in this manner. This contrasts with previous studies in cultured cells, where mNG2(Δ11) was transfected or virally transduced (Cabantous et al., 2005; Kamiyama et al., 2016; Leonetti et al., 2016), presumably resulting expression of multiple copies and greater apparent sensitivity. By contrast, in the GOGREEN system, *mNG2(Δ11)* is expressed from a single genomic locus, and therefore copy number is more limited than it may be in cell culture systems relying on random integration followed by clonal selection for the brightest cell line. Nevertheless, we find that the GOGREEN system is actionable for abundant proteins, opening new opportunities for imaging many proteins in vivo.

Curiously, a *GFP* knock-in reporter of *Nanog* expression (Maherali et al., 2007) is sufficiently bright so as to be detected in embryos (Frum et al., 2019). Yet, we were unable to detect *Nanog* using split mNG2, in spite of the fact that mNG is reportedly up to three times brighter than GFP (Shaner et al., 2013). We therefore caution that splitting and reconstitution could diminish overall fluorescence, when compared with full-length proteins. Consistent with this proposal a prior study found split mNG2 to be half as bright as the full-length mNG protein from which it was derived (Feng et al., 2017). For all these reasons, we recommend use of the V5 tagging system where visualization of proteins of lower abundance is the priority.

One caveat with analyzing F0 knock-in embryos is the potential for mutagenesis during the knock-in. For this reason, we preferentially targeted the protein tags to the C-terminal region, so as to avoid null allele-producing indels due to CRISPR/Cas9-mediated double-stranded DNA breaks. Thus, it is important to take this into consideration when implementing the GOGREEN or V5 systems. An additional caveat to be considered when planning to analyze F0 knock-in embryos is the potential for genetic mosaicism. The degree of genetic mosaicism was observed to be minimal in our studies. This a limitation that can be addressed by experimental replication to provide an understanding of a particular gene’s expression pattern that is more representative. At the very least, the GOGREEN and V5 systems provide preliminary insight into a given gene’s expression pattern that can be helpful in guiding future experiments (e.g., whether to screen for protein-specific antibodies, whether to establish a permanent mouse lines, or whether to make a knock-out mouse model).

### Opportunities

The GOGREEN and V5 systems present exciting opportunities for biological investigation outside of preimplantation mouse development as well. For example, V5 or mNG2(11) knock-in embryos could be transferred to recipient females to allow for postimplantation development so that protein localization can be evaluated in later developmental processes. Additionally, transferred knock-in embryos could be allowed to gestate to establish mouse lines for subsequent imaging of protein localization in adult tissues and processes. Alternatively, both the GOGREEN and V5 systems are amenable to somatic cell introduction, for example via viral transduction (Yoon et al., 2018). Our studies thus provide guidelines, molecular reagents, and genotyping assays to enable these applications.

We note opportunities for applying biochemical and molecular techniques in vivo as well. V5 is commonly used for purifying proteins from cells and tissues for the downstream identification of protein or nucleotide interactions, including immunoprecipitation-western blotting or mass spectrometry, chromatin-immunoprecipitation (ChiP) and quantitative reverse transcription polymerase chain reaction (qPCR) or sequencing or ribonucleotide pulldown and sequencing (RIP-seq). We therefore envision that the approaches described here could be used to generate stable mouse lines that enable anti-V5 antibody-mediated discovery of protein localization patterns, protein and RNA binding partners and transcription factor DNA binding sites and sequences.

Finally, both the GOGREEN and V5 systems could also be used in other animals, where breeding to establish knock-in lines is either impractical or inappropriate, including humans or other primates. There would be additional advantages to applying either system to emerging mammalian models, such as marsupials, where protein-specific antibodies have not yet been developed. For live imaging, *mNG2(Δ11)* could be provided by mRNA injection, while *mNG2(11)* would be knocked in frame into genes of interest. If fixed imaging of low abundance proteins is preferred, then V5 could be knocked in. Either system promises new opportunities for discovery of developmental principles in mouse as well as understudied mammalian species.

## Acknowledgements

We thank current members of the Ralston Lab as well as Dr. Tristan Frum for helpful discussion and training, Axel Schmitter for assistance in the lab, and Drs. Elena Demeriva and Huirong Xie at the MSU Transgenic and Genome Editing Facility for the *R26* mouse knock-in line. Research in the Ralston Lab is supported by NIH R35 GM131759 to A.R.

## Author Contributions

A.R. conceptualized the study and acquired funding. A.R. and D.O. designed the experiments, D.O. performed and interpreted the experiments. A.R. and D.O. wrote the paper.

---

**Figure S1.**
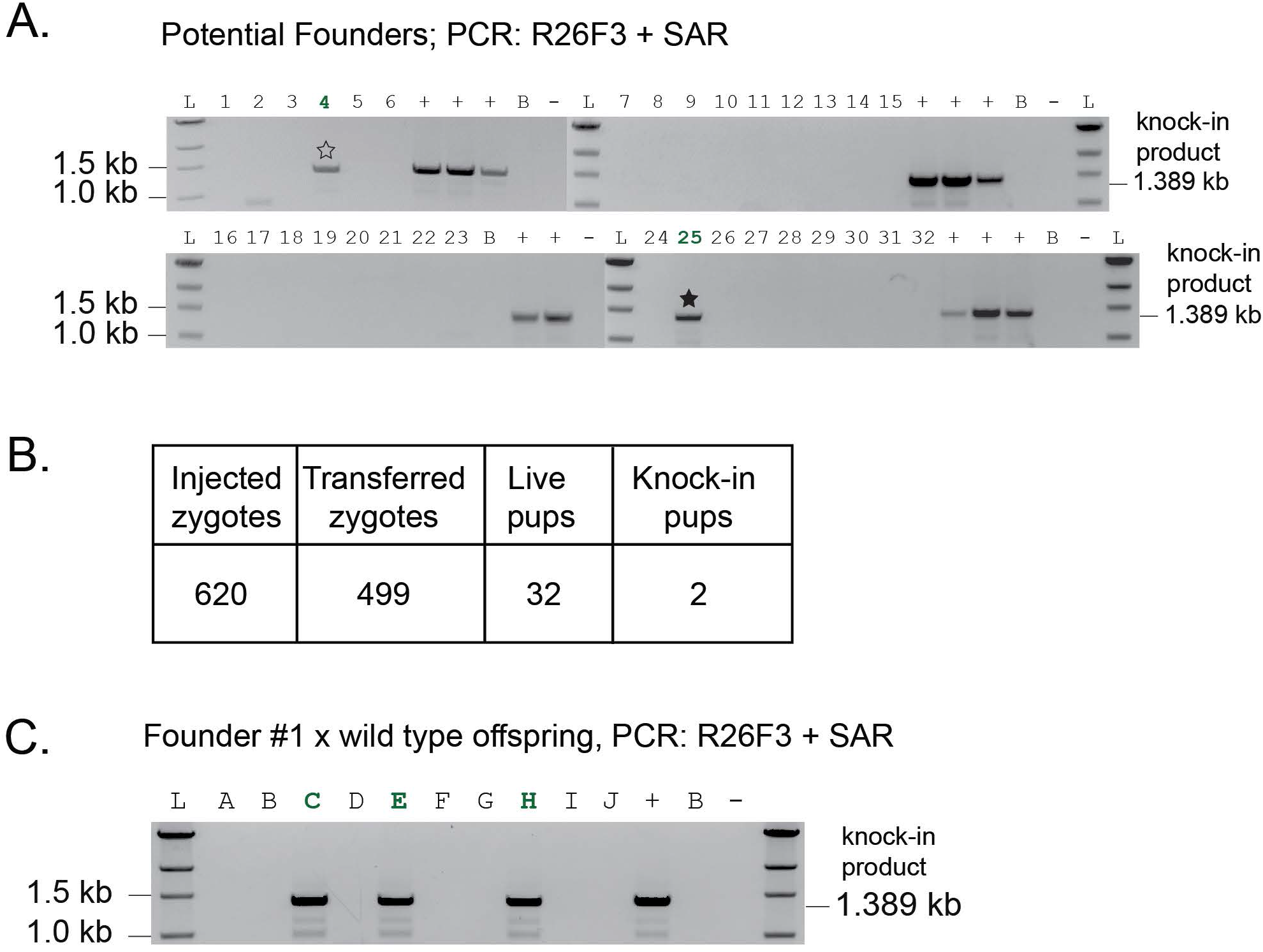
Details of the *R26-mNG2(Δ11) mouse line*. A) PCR genotyping of tail tip biopsies from offspring born following zygote injection of CRISPR/Cas9 reagents to target the *CAG-mNG2(Δ11)* expression construct to the *Rosa26* locus. Injected zygotes were subsequently transferred to recipient mothers to enable gestation and birth of potential founders. Successful homologous recombination suggested by amplification of 1.389 kb band by PCR from genomic DNA. L = DNA ladder, numbers indicate individual mice screened by PCR genotyping, + = positive control (targeted ES cells), - = negative control (no DNA template), B = C57BL/6 wild type genomic DNA, stars = potential founders. B) Summary of CRISPR/Cas9 *R26* targeting with *mNG2(Δ11)* expression construct. C) PCR genotyping of F1 offspring resulting from breeding of a potential founders (#1, hollow star in panel A) to wild type mouse. Detection of 1.389 kb band by PCR genotyping of genomic F1 DNA confirms germline transmission of the *R26-mNG2(Δ11)* allele.

**Figure S2.**
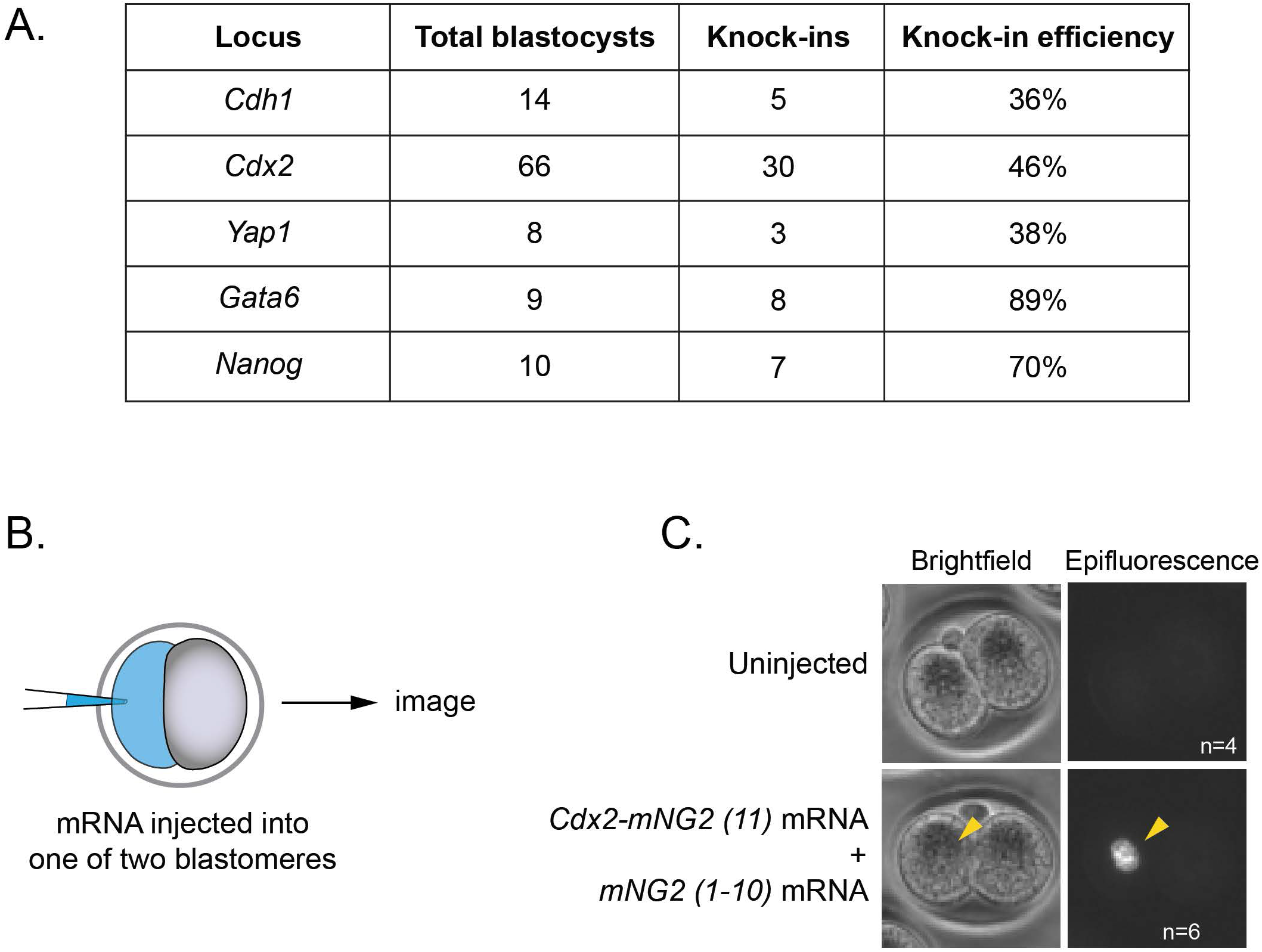
Overexpression of components of the GOGREEN system can enable detection of low abundance nuclear proteins. A) Summary of *mNG2(11)* knock-ins, as determined by PCR genotyping, that were undetectable using the GOGREEN system. B) Overview of experimental design: mRNA injection into one of two blastomeres of the early mouse embryo, followed by imaging. C) Overexpression of mRNAs encoding *mNG2(Δ11)* and *Cdx2-mNG2(11)* leads to reconstituted fluorescence in the nucleus of the injected blastomere. n = number of embryos presenting the phenotype shown.

## Methods

### Animals

All animal research was conducted in accordance with the guidelines of the Michigan State University Institutional Animal Care and Use Committee. CD-1 mice were kept on a 12-hour day/night cycle and fed ad libitum.

### Plasmid construction

The pR26-CAG-mNG2(Δ11) expression/targeting vectors was cloned by insertion of an IDT gBlock encoding mNG2(Δ11) into the previously published vector pR26-AsisI/MluI (Addgene #74286) (Chu et al., 2016). After cloning, the Lox-Stop-Lox site was removed by exposure to recombinant Cre recombinase (NEB), using the NEB standard protocol. The expression vector pCAG-mNG2(11) were derived from pCAG-ERT2CreERT2 (Addgene #13777) (Matsuda and Cepko, 2007) by replacing the coding sequence with an IDT gBlock (Table S4) encoding mNG2(11) via restriction ligation with EcoRI and NotI. The in-vitro transcription plasmids for *Clta-mNG2(11), mNG2(Δ11)*, and *Cdx2-mNG2(11)* were cloned by inserting an IDT gBlock (Table S4) containing the respective coding sequences into a pcDNA3.1-poly(A)83 vector (Yamagata et al., 2005) downstream of the T7 promoter via restriction ligation with HindIII and NotI. PX459-sgRosa26-1 was generated by inserting the guide RNA sequence targeting *Rosa26* (Table S3) into pSpCas9(BB)-2A-Puro (PX459) V2.0 (Addgene) via restriction ligation with BbsI.

### In vitro transcription

In vitro transcription was performed using the T7 mMessage mMachine kit (Life Technologies). Each IVT construct was digested with XbaI, followed by ethanol precipitation, and was then used in the IVT reaction per manufacturer’s instructions. Resulting mRNA was purified using the MEGAclear kit (Ambion). mRNA quantity and quality were assessed by Nanodrop spectrophotometer and by agarose gel, respectively.

### Zygote and 2-cell embryo microinjection

Target-specific crRNA and non-variable tracrRNA obtained from IDT were each suspended in injection buffer (1 mM Tris HCl, pH 7.5; 0.1 mM EDTA), mixed at a 1:1 molar ratio and annealed in a thermal cycler by ramping down from 95 °C to 25 °C at 0.1 °C/second. The annealed RNAs were then mixed with recombinant Cas9 Protein at a 5:1 RNA:Cas9 molar ratio and allowed to form RNPs for 15 minutes at room temperature. RNPs were mixed with donor ssODN synthesized as Ultramers from IDT (Table S2) and diluted to working concentrations (Table S5) by addition of injection buffer. For mRNA injections, each mRNA was diluted in injection buffer to 350 ng/µl and injected either into one pronucleus of the mouse zygote or into one blastomere of the 2-cell mouse embryo. Injection mixes and mRNAs were aliquoted and stored at -80 °C, avoiding freeze/thaw cycles.

Zygotes were harvested from naturally mated pregnant mice on the day that copulatory plugs were detected. Oviducts were flushed with M2 medium (Millipore Sigma), and then injection mix was delivered into one pronucleus or the nucleus of one blastomere via microinjection (Nagy et al., 2003). Injected zygotes were cultured in KSOM + AA (Millipore Sigma) for up to five days before being either fixed or imaged live. Only embryos that survived injection and appeared to have cavitated or to be attempting cavitation were included in the analysis.

### Generation of R26-mNG2(Δ11) Mouse Line

The *R26-mNG2(Δ11)* mouse line was generated by zygote microinjection at the MSU Transgenic and Genome Editing Facility. A mixture containing 5 ng/µl circular pR26-CAG-mNG(Δ11) and 125 ng/µl Cas9 RNP complexed with *sgRosa26* guide RNA was injected into one pronucleus of C57BL/6J mouse zygotes. Zygotes were then transferred to CD-1 recipient mice. After birth, tail tips were screened by PCR for successful integration of the *R26-CAG-mNG(Δ11)* allele.

### Immunofluorescence and Confocal Microscopy

V5 was detected by mouse anti-V5 antibody (Invitrogen). Embryos were fixed with 4% formaldehyde (Polysciences) for 10 min, permeabilized with 0.5% Triton X-100 (Millipore Sigma) for 30 min, and then blocked in 10% Fetal Bovine Serum (Hyclone) with 0.1% Triton X-100 for 1 hr at room temperature. Embryos were then incubated in V5 at a dilution of 1:400 in blocking solution at 4 °C overnight. The next day, embryos were stained with goat anti-mouse Alexa488 (Invitrogen) at a 1:400 dilution in blocking solution for 1 hour at room temperature. Embryos were then stained for 10 min at room temperature in 50 µM Hoechst nucleic acid stain (Thermo Fisher). Split mNG2 embryos were imaged either fixed or live after Hoechst staining. Imaging was performed using an Olympus FluoView FV1000 Confocal Laser Scanning Microscope system with 20x UPlanFLN objective (0.5 NA) and 5x digital zoom. For each embryo, z-stacks were collected with 3 µm intervals between optical sections. Optical sections are displayed as an intensity projection over the Z axis.

### Genotyping

To genotype embryos, genomic DNA was extracted from single blastocysts by placing each blastocyst in a microtube containing 4.4 µl Extraction buffer (REDExtract-N-Amp Tissue PCR Kit, Millipore Sigma) mixed with 1.1 µl of Tissue Prep Buffer, and then incubating tubes at 56 °C for 30 minutes, 24 C for 5 minutes, and 95 °C for 5 minutes. After incubation, 5 µl Neutralization buffer was added to each tube. In subsequent reactions, 1 µl of embryo extract was used as PCR template, and locus-specific primers (Table S1).

To genotype adult mice, genomic DNA was extracted from ear punch biopsies using the Wizard SV Genomic DNA Purification System (Promega), and PCR was performed using Herculase II Polymerase (Agilent).

### Sequencing

To confirm the identity of select PCR products, the products were directly cloned into pCR2.1 TOPO using the Invitrogen TOPO TA Cloning Kit (Invitrogen). Plasmids containing the PCR product were prepped with the Spin Miniprep Kit (Qiagen) and sent to the RTSF Genomics Core at Michigan State University for Sanger sequencing.

### mNG2(Δ11) ES Cell Derivation

R1/E ES cells (ATCC) were cultured on CF-1 feeder MEFs (Applied Stem Cell) in ES cell medium [DMEM supplemented with 1000 U/mL leukemia inhibitory factor (Millipore Sigma), 15% (v/v) fetal bovine serum (Hyclone), 2 mM L-Glutamax (Thermo Fisher), 0.1 mM beta-mercaptoethanol (Millipore Sigma), 0.1 MEM non-essential amino acids (Millipore Sigma), 1 mM Sodium Pyruvate (Millipore Sigma), and 1% (v/v) penicillin/streptomycin (Gibco)]. Passage 13 R1/E ES cells were cultured to approximately 70% confluence in a 10 cm dish, and then electroporated with pX459-sgRosa26-1 and pR26-CAG-mNG2(Δ11) as follows: pelleted cells were resuspended in 800 µl Embryo-Max Electroporation Buffer (Millipore Sigma) containing 20 µg each plasmid, and cells were then electroporated in a 0.4 cm electrode gap electroporation cuvette (Bio-Rad) using Bio-Rad Gene Pulser XCell electoporator (250 V, 500 µF, infinite Ω). Subsequently, 400 µl electroporated cells were then diluted in 10 mL ES cell medium, and then plated on a 10 cm dish on puromycin-resistant DR4 feeder MEFs (Applied Stem Cell). After 24 hours, selection was started with ES cell medium containing 1.25 µg/mL puromycin (Gibco). After 12 days, colonies were picked into 96-well plates and expanded over several more passages. Cell lines were genotyped by PCR using R26F3 and SAR primers to detect insertion of *mNG2(Δ11)* in the *Rosa26* locus (Table S1).

## DATA AND CODE AVAILABILITY

This study did not generate any data sets or code.

---

**Supplementary Table 1.**
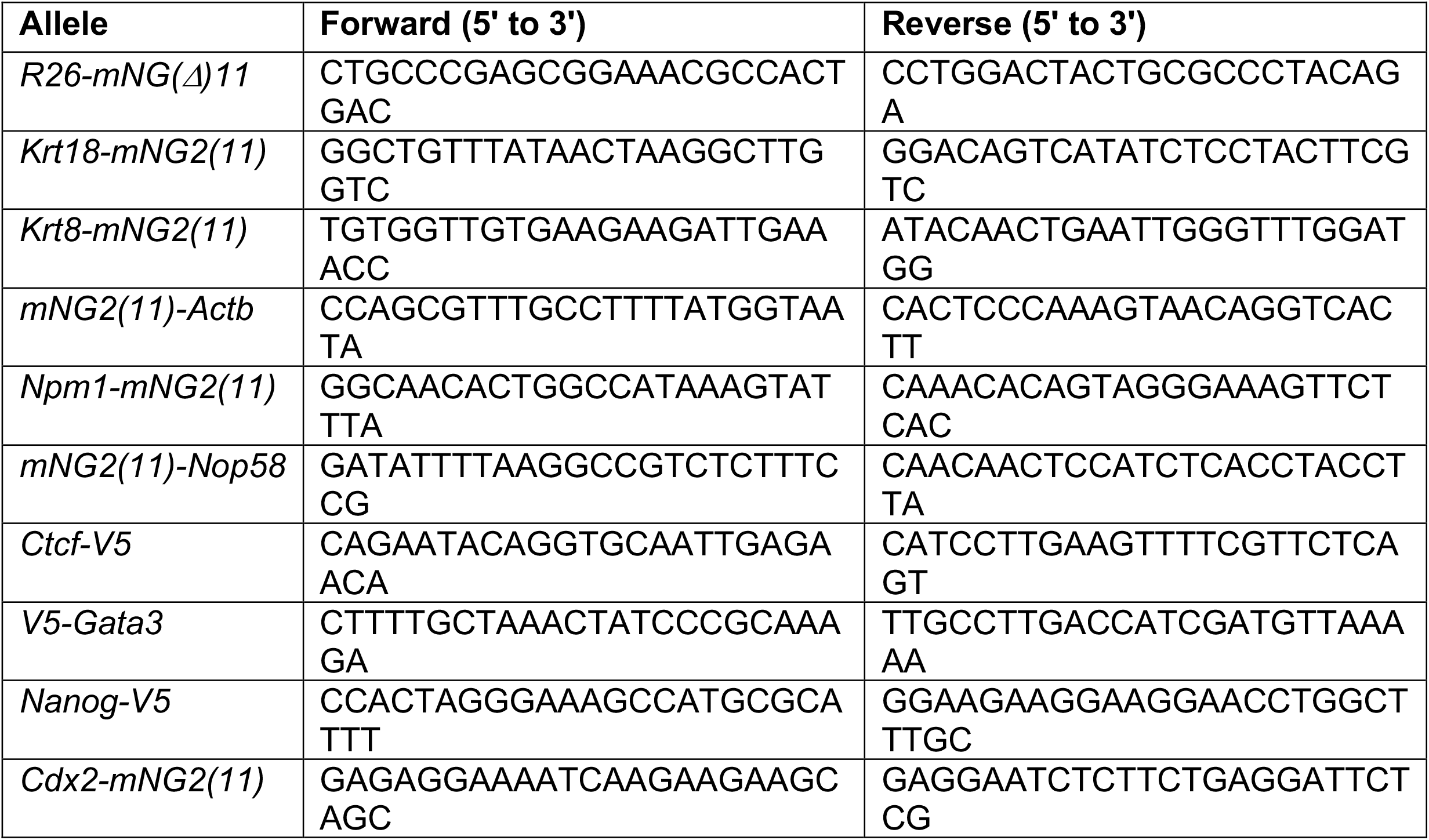
Primers used in this study.

**Supplementary Table 2.**
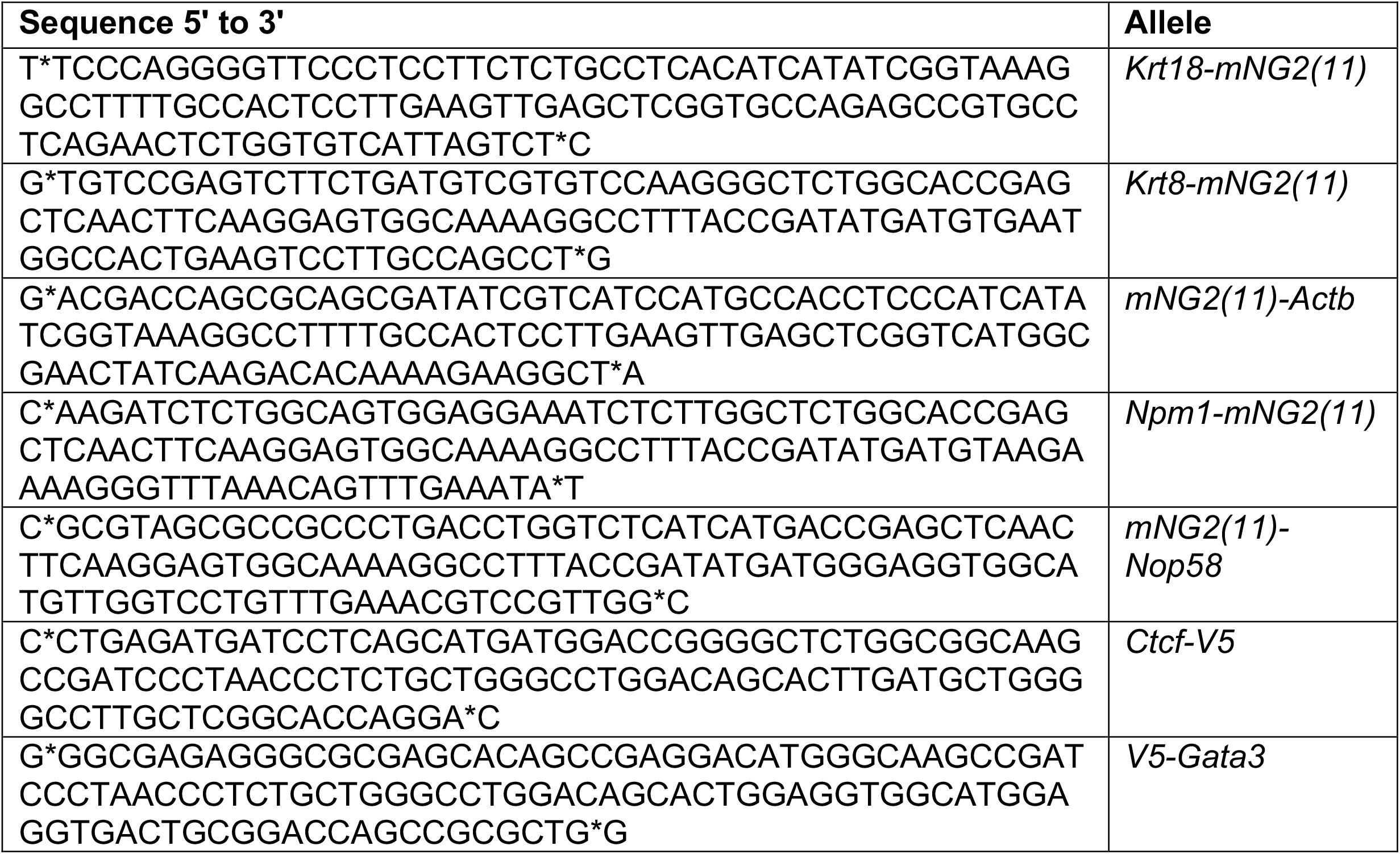
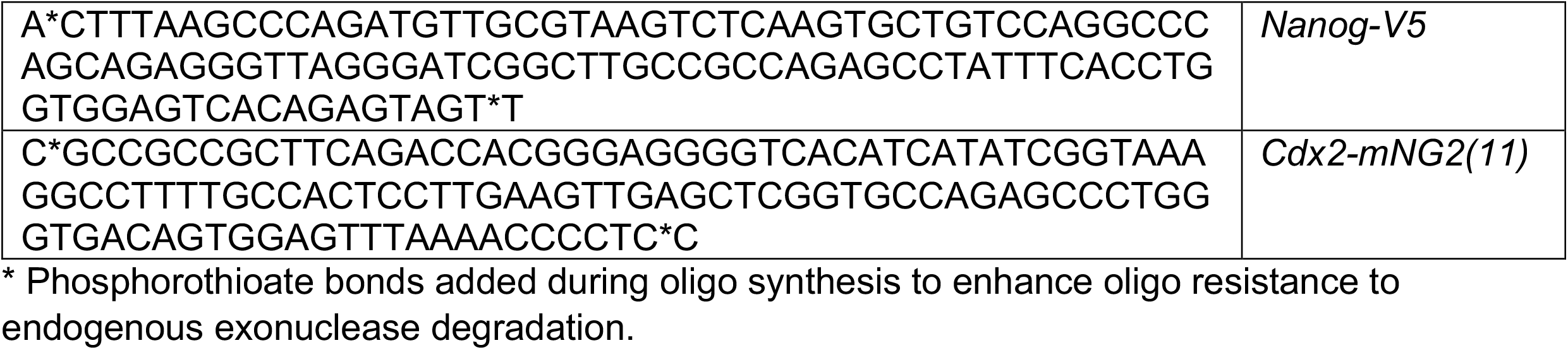
Synthesized ssOND sequences used in this study

**Supplementary Table 3.**
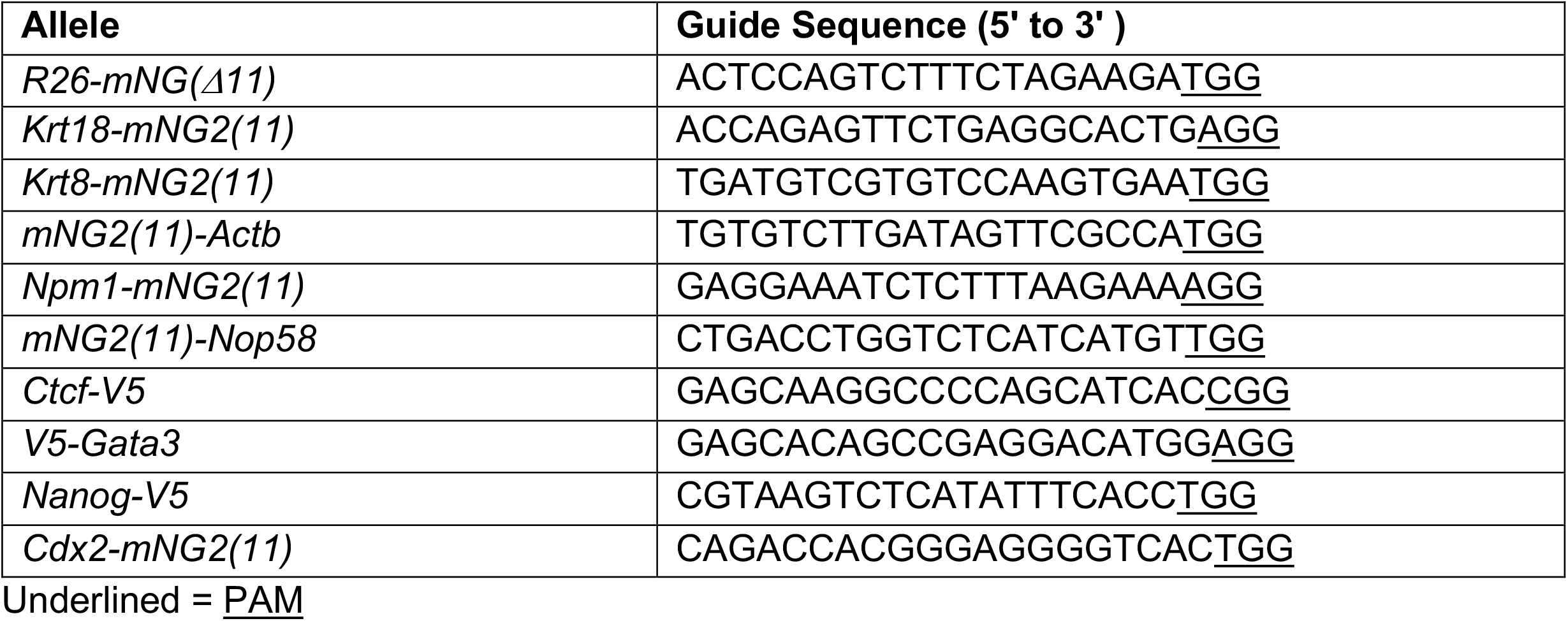
CRISPR Guides used in this study

**Supplementary Table 4.**
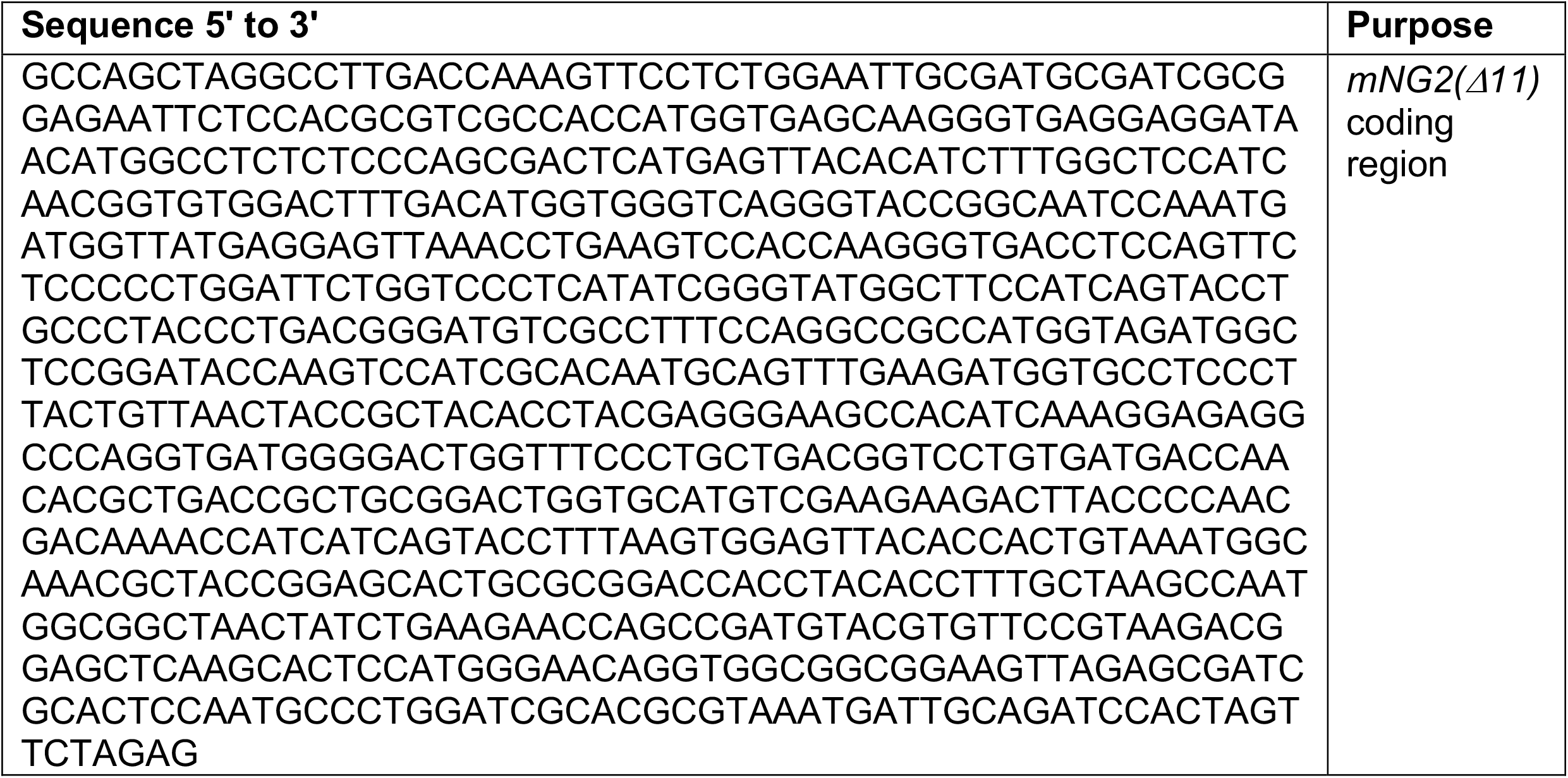
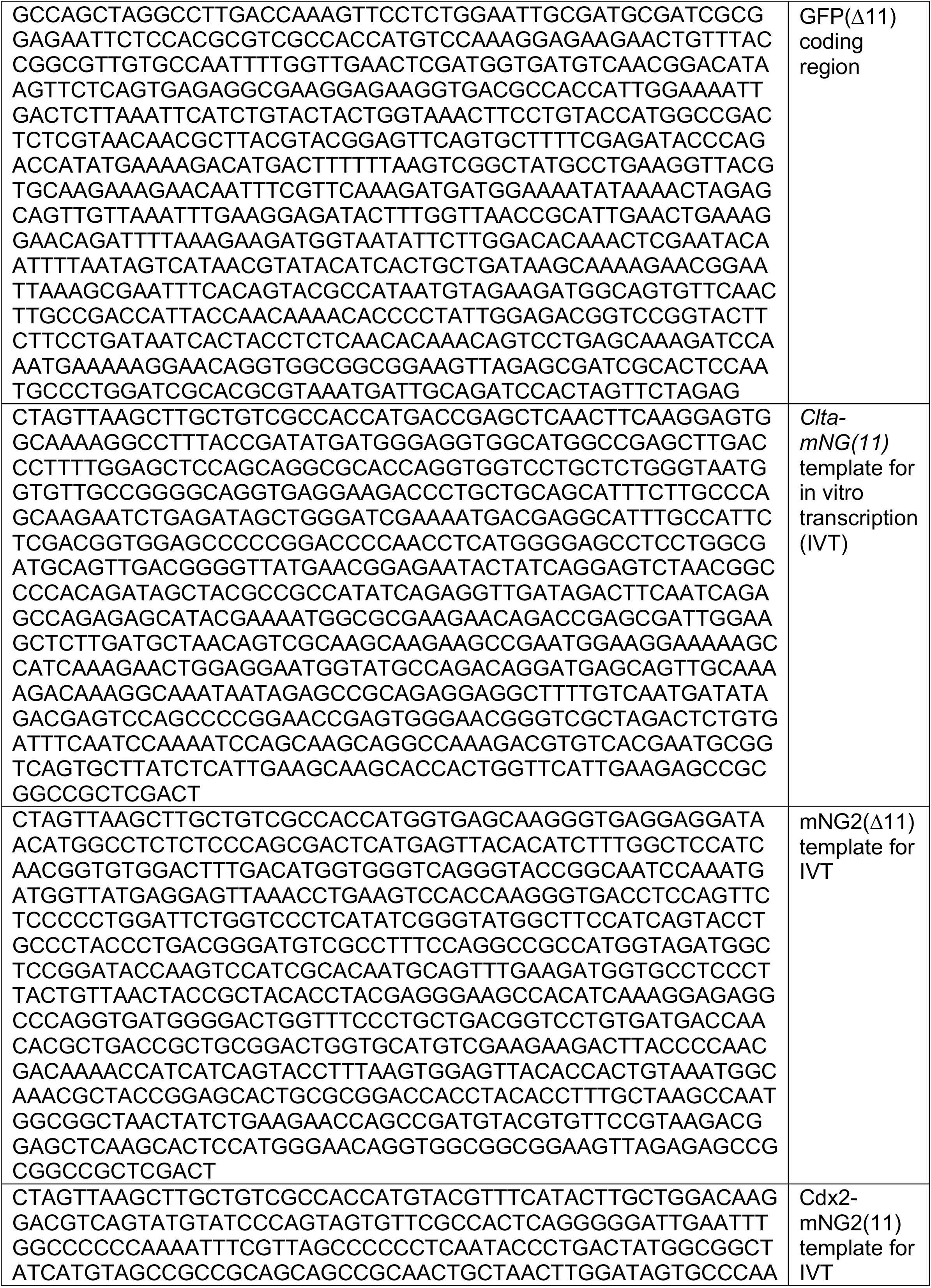
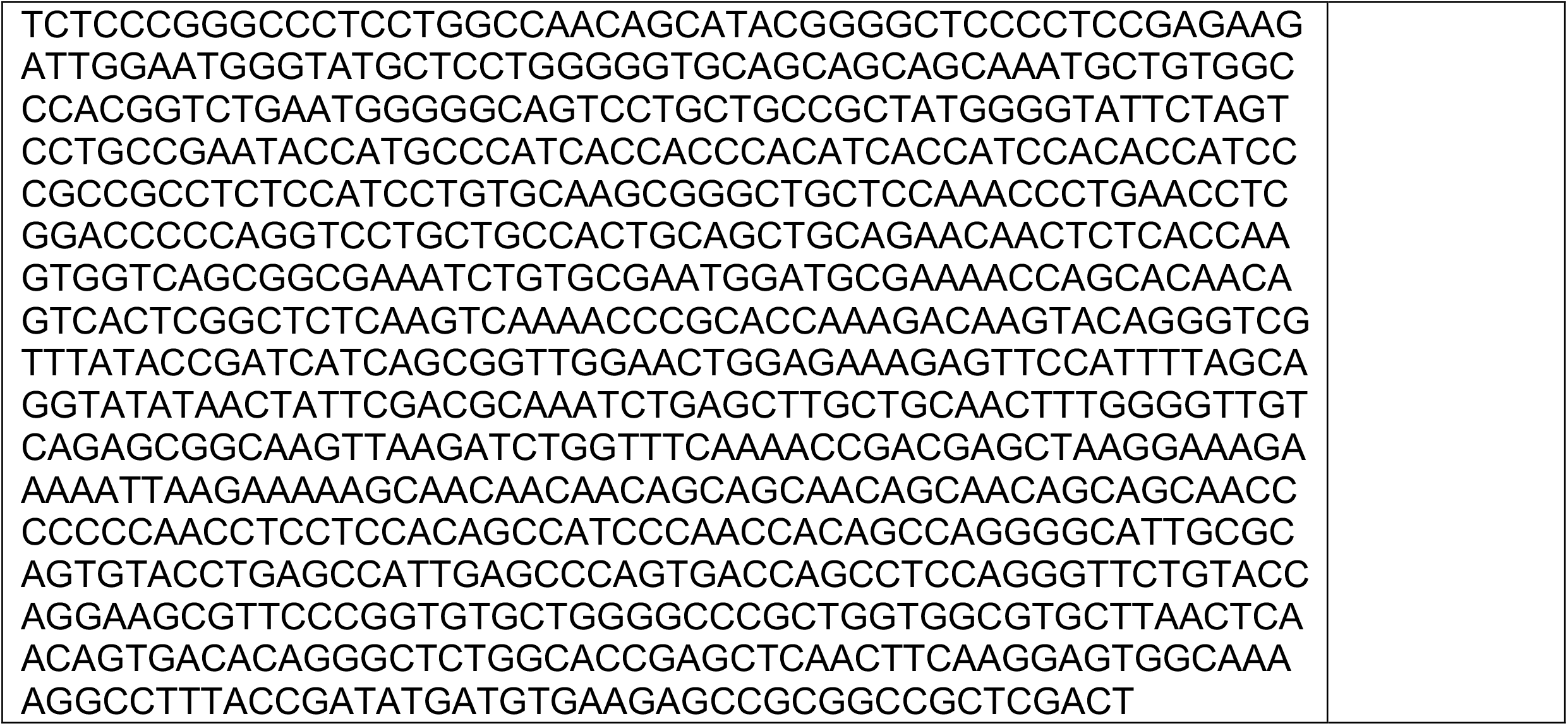
Other oligos synthesized for this study

**Supplementary Table 5:** Concentrations of Cas9 RNP and single stranded oligonucleotide donor (ssOND) used for targeting each gene.

| Gene Targeted | Tag | Cas9 RNP (ng/μl) | ssOND (ng/μl) |
| --- | --- | --- | --- |
| *Krt8* | mNG2(11) | 100 | 20 |
| *Actb* | mNG2(11) | 100 | 20 |
| *Krt18* | mNG2(11) | 100 | 20 |
| *Nop58* | mNG2(11) | 100 | 20 |
| *Npm1* | mNG2(11) | 100 | 10 |
| *Ctcf* | V5 | 25 | 5 |
| *Gata3* | V5 | 100 | 20 |
| *Nanog* | V5 | 100 | 20 |

